# Molecular architecture determines brain delivery of a transferrin-receptor targeted lysosomal enzyme

**DOI:** 10.1101/2021.05.21.445035

**Authors:** Annie Arguello, Cathal S. Mahon, Meredith E. K. Calvert, Darren Chan, Jason C. Dugas, Michelle E. Pizzo, Elliot R. Thomsen, Roni Chau, Lorna A. Damo, Joseph Duque, Timothy Earr, Meng Fang, Tina Giese, Do Jin Kim, Nicholas Liang, Isabel A. Lopez, Hoang N. Nguyen, Hilda Solanoy, Buyankhishig Tsogtbaatar, Julie C. Ullman, Junhua Wang, Mark S. Dennis, Dolores Diaz, Kannan Gunasekaran, Kirk R. Henne, Joseph W. Lewcock, Pascal E. Sanchez, Matthew D. Troyer, Jeffrey M. Harris, Kimberly Scearce-Levie, Lu Shan, Ryan J. Watts, Robert G. Thorne, Anastasia G. Henry, Mihalis S. Kariolis

## Abstract

Delivery of biotherapeutics across the blood-brain barrier (BBB) is a challenge. Many approaches fuse biotherapeutics to platforms that bind the transferrin receptor (TfR), a brain endothelial cell target, to facilitate receptor-mediated transcytosis across the BBB. Here, we characterized the pharmacological behavior of two distinct TfR-targeted platforms fused to iduronate 2-sulfatase (IDS), a lysosomal enzyme deficient in mucopolysaccharidosis type II (MPS II), and compared the relative brain exposures and functional activities of both approaches in mouse models. IDS fused to a moderate-affinity, monovalent TfR binding enzyme transport vehicle (ETV:IDS) resulted in widespread brain exposure, internalization by parenchymal cells, and significant substrate reduction in the CNS of an MPS II mouse model. In contrast, IDS fused to a standard high-affinity bivalent antibody (IgG:IDS) resulted in lower brain uptake, limited biodistribution beyond brain endothelial cells, and reduced brain substrate reduction. These results highlight important features likely to impact the clinical development of TfR-targeting platforms in MPS II and potentially other CNS diseases.

**Summary:** Brain delivery, biodistribution and pharmacodynamics of a lysosomal enzyme fused to a moderate-affinity transferrin receptor-directed blood-brain barrier enzyme transport vehicle are superior to a traditional high-affinity anti-TfR monoclonal antibody fusion.

## Introduction

The use of protein-based therapies to treat neurodegenerative diseases has been limited by minimal brain exposure following systemic administration (Kumar et al., 2018; Stanimirovic et al., 2018). Most polar small molecules and nearly all macromolecules are effectively restricted from reaching the brain in therapeutically relevant concentrations by physical and biochemical barriers, most notably the blood-brain barrier (BBB) (Abbott et al., 2018; Banks, 2016). Brain endothelial cells that form the BBB have several unique physiological properties that distinguish them from peripheral endothelial cells, including tight junctions, relatively low endocytic activity, and the expression of numerous transporters and receptors (Profaci et al., 2020). As a result, CNS concentrations of antibodies often reach only about 0.01%-0.1% (Atwal et al., 2011; Poduslo et al., 1994; St-Amour et al., 2013) of peripheral levels after systemic administration and, typically, much of the brain-associated antibody is confined to the endothelium and not parenchymal cells (St-Amour et al., 2013).

A promising strategy to improve brain uptake of biotherapeutics leverages receptor-mediated transcytosis (RMT) at the BBB (Banks, 2016; Lajoie and Shusta, 2015). RMT is an endogenous process wherein essential biomolecules that cannot passively diffuse into the brain from the bloodstream are actively transported across brain endothelial cells via specific receptors on their luminal surface (Johnsen et al., 2019). While some brain endothelial cell receptors capable of initiating RMT are downregulated postnatally (e.g. mannose-6-phosphate receptor) (Urayama et al., 2004; Urayama et al., 2008), other receptors capable of RMT such as the transferrin receptor (TfR) are expressed throughout life (Preston et al., 2014). Biotherapeutic platforms engineered to interact with persistently expressed receptors can therefore exploit RMT pathways to gain access to the CNS (Johnsen et al., 2019; Jones and Shusta, 2007).

TfR has been among the most studied RMT targets at the BBB (Johnsen et al., 2019), owing in part to its enriched expression on brain endothelial cells (Jefferies et al., 1984) and its constitutive ligand-independent endocytosis (Hopkins et al., 1985). Many platforms targeting TfR have been described (Terstappen et al., 2021), including conventional high-affinity bivalent antibodies (Friden et al., 1991), bispecific antibodies (Yu et al., 2011), antibody fragments (Lesley et al., 1989), peptides (Kuang et al., 2016), antibody-fusion architectures (Hultqvist et al., 2017; Niewoehner et al., 2014; Sonoda et al., 2018) and, most recently, a transport vehicle (TV) consisting of an Fc domain engineered to directly bind TfR (Kariolis et al., 2020). Of these, traditional antibodies directed against target receptors have several attractive features, most notably, established discovery and development methods to generate specificity and high affinity. Several antibodies have been reported that engage TfR bivalently with sub-nanomolar apparent affinities (Pardridge, 2015; Sonoda et al., 2018). While such antibodies are capable of being internalized into brain endothelial cells, a number of imaging and biodistribution studies in mouse models have suggested they may be only minimally released into the brain parenchyma (Moos and Morgan, 1998; Paris-Robidas et al., 2011; Paterson and Webster, 2016; Yu et al., 2011). Studies utilizing monovalent anti-TfR antibodies with weaker affinity, have shown enhanced BBB transcytosis and brain accumulation (Bien-Ly et al., 2014; Weber et al., 2018; Yu et al., 2011). A proposed mechanism for the increased brain uptake is altered cellular trafficking, whereby weaker affinity anti-TfR antibodies avoid sorting to lysosomes and subsequent degradation, while high-affinity anti-TfR antibodies mainly accumulate in lysosomes driving receptor degradation (Bien-Ly et al., 2014). Several studies have demonstrated that the intrinsic properties of TfR-directed architectures (including affinity and valency) can impact transport across the BBB (Dennis and Watts, 2012; Moos and Morgan, 2001; Villasenor et al., 2019; Villasenor et al., 2017; Yu et al., 2011). Despite these studies, questions around the most suitable TfR-targeting molecular architecture for optimal brain delivery have remained. This is particularly relevant for lysosomal storage disorders (LSDs), where traditional high-affinity, bivalent TfR-binding antibody fusions and newer monovalent TfR-binding TV-fusions are currently being evaluated in the clinic (NCT04251026; Okuyama et al., 2019).

The primary treatment for LSDs involves ERTs that have limited transport across the BBB and therefore represent an attractive candidate cargo to examine the relative merits of specific TfR-based approaches. LSDs represent a family of more than 50 monogenic diseases, many of which are characterized by a defect in a single lysosomal enzyme (Neufeld, 1991; Schultz et al., 2011). Disease-associated variants lead to a reduction or loss of enzymatic activity, resulting in substrate accumulation and broad lysosomal dysfunction (Platt et al., 2012). Perturbed lysosomal function can trigger pathogenic cascades affecting multiple tissues throughout the body, including the CNS (Bellettato and Scarpa, 2010). At present, the standard of care for many LSDs is systemically administered recombinant ERTs; however, these enzymes do not readily cross the BBB and typically have been ineffective in treating the CNS manifestations of disease (Desnick and Schuchman, 2012; Muldoon et al., 2013; Scarpa et al., 2015). Mucopolysaccharidosis II (MPS II) is an X-linked LSD resulting from deficient activity of iduronate-2-sulfatase (Schultz et al.), an enzyme responsible for the catabolism of the glycosaminoglycans (GAGs) heparan and dermatan sulfate (Wraith et al., 2008). MPS II is characterized by widespread GAG accumulation with a host of secondary pathologies and nearly 70% of patients present with neuronopathic disease (Noh and Lee, 2014). Since its approval in 2006, recombinant IDS has transformed the clinical management of MPS II, successfully reducing GAG accumulation in the periphery (Muenzer et al., 2006; Sohn et al., 2013). However, administration of recombinant IDS does not effectively treat CNS pathology (Parini and Deodato, 2020; Scarpa et al., 2017), highlighting the critical need for the development of new brain-penetrant therapies for MPS II.

We recently described a TV-based biotherapeutic for MPS II generated by fusing IDS to an engineered TfR-binding Fc fragment (ETV:IDS) (Kariolis et al., 2020; Ullman et al., 2020). ETV:IDS binds TfR monovalently with a moderate affinity identified to maximize brain uptake, in contrast to most other TfR-based enzyme platforms where enzymes have been fused to high-affinity, bivalent anti-TfR antibodies (Boado et al., 2018; Sonoda et al., 2018; Zhou et al., 2012). Characterizing how these different molecular architectures impact CNS biodistribution and potency therefore has significant relevance for their clinical translation.

Here, we compare ETV:IDS to a high-affinity, bivalent TfR-binding antibody-enzyme fusion with IDS (IgG:IDS) in order to determine which format most effectively enables broad biodistribution of IDS to the brain following systemic administration. Biodistribution was quantitatively assessed using a combination of bulk tissue measurements, tissue fractionation, fluorescence imaging and super resolution confocal microscopy. Activity was evaluated by measurements of total GAG levels in liver, brain, and cerebrospinal fluid (CSF) in a mouse model of MPS II. We demonstrated that ETV:IDS results in enhanced brain exposure compared to IgG:IDS. Importantly, ETV:IDS reduced brain and CSF GAG levels to a greater extent than IgG:IDS likely as a direct reflection of improved biodistribution.

## Results and Discussion

### Biochemical characterization of ETV:IDS and IgG:IDS demonstrates impact of architecture on receptor affinity

ETV:IDS was engineered by fusing IDS to the N-terminus of the TV, as previously described (Ullman et al., 2020), while IgG:IDS was generated by fusing IDS to the C-terminus of both heavy chains of a high-affinity, anti-TfR antibody (Fig. 1A). IDS requires several post-translational modifications (PTMs) for proper biological function (Demydchuk et al., 2017; Millat et al., 1997) that can be altered by the expression and purification processes. In order to distinguish parameters created by process (e.g. PTMs) from properties intrinsic to platform architecture (e.g. affinity, valency), we first characterized the *in vitro* biochemical enzymatic activity of purified ETV:IDS and IgG:IDS. As ETV:IDS production and associated activities have been previously described (Ullman et al., 2020) we focused here on generating active IgG:IDS. Variations in the expression conditions yielded batches of IgG:IDS with increasing levels of specific activity (benchmarked to idursulfase; Fig. 1B left panel IgG:IDS #1-3). Using optimized conditions, a batch of IgG:IDS which had 1.5-fold higher IDS activity compared to ETV:IDS (Fig. 1B right panel) was generated for further characterization as the most stringent comparison against ETV:IDS in biodistribution and efficacy studies.

**Figure 1.**
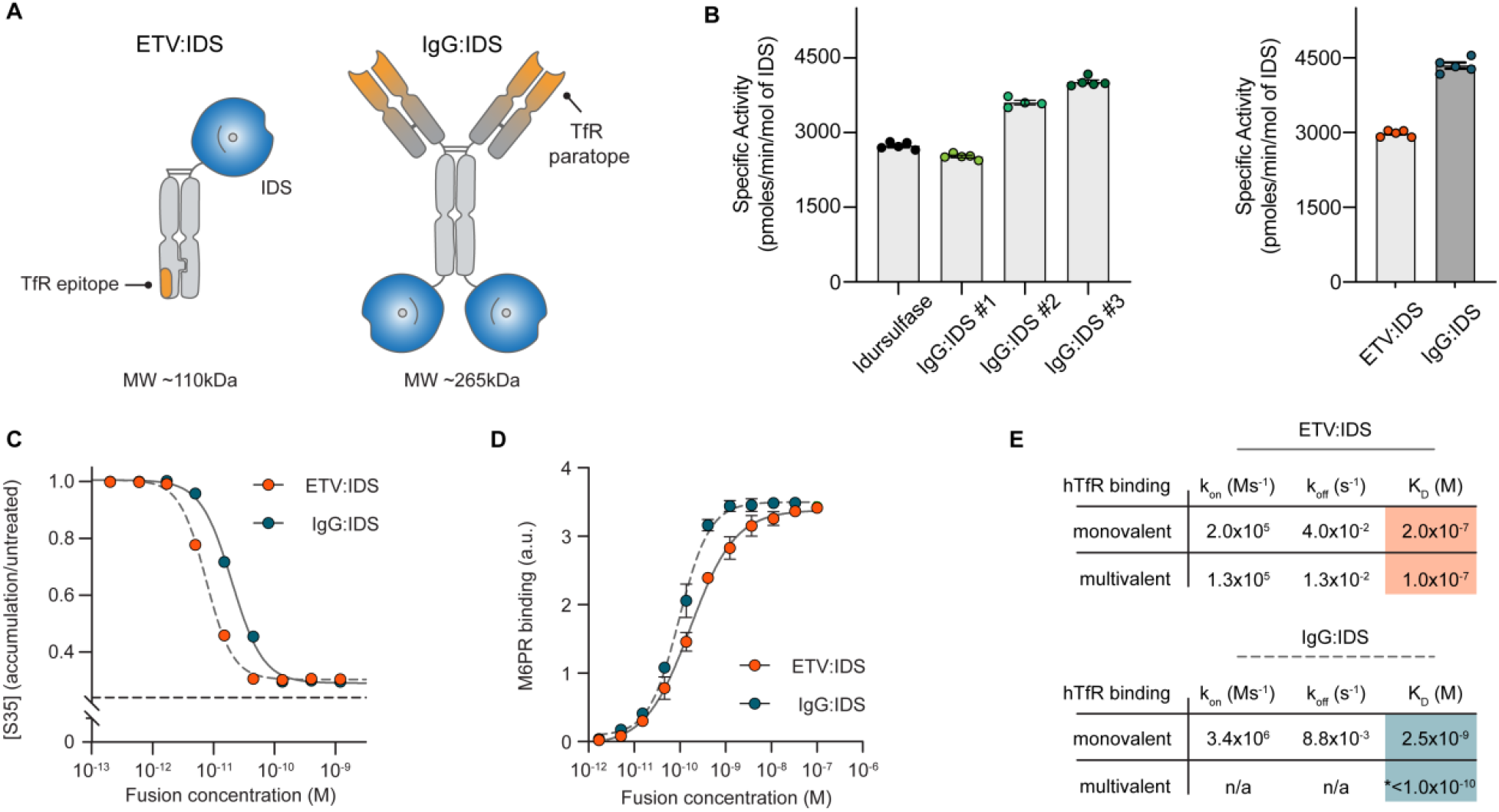
Biochemical characterization of ETV:IDS and IgG:IDS. **(A)** ETV:IDS is a fusion of the lysosomal enzyme iduronate 2-sulfatase (IDS) to the Transport Vehicle, a TfR-binding Fc domain. IgG:IDS is a high affinity anti-TfR huIgG fused to IDS at the C- terminus of each heavy chain. **(B)** Specific activities of ETV:IDS and IgG:IDS were measured using a synthetic fluorogenic substrate. Graphs display mean ± SD. Samples represented: Left panel: Idursulfase (commercially approved recombinant IDS), three IgG:IDS preparations generated by co-transfecting CHO cells with increasing amounts of SUMF1 leading to increasing specific activity; Right panel: ETV:IDS and high activity IgG:IDS were chosen for further characterization. (**C**) ^35^S sulfate-labeled substrates in MPS II patient derived fibroblasts after treatment with ETV:IDS or IgG:IDS. n = 3 experiments with three patient lines per phenotype used in each experiment. Dashed line represents the amount of ^35^S-labeled substrate in healthy control cells. (**D**) Binding affinities of ETV:IDS and IgG:IDS to the mannose-6-phosphate receptor (M6PR) were determined by ELISA; n = 3 technical replicates, representative graph shown. **(E)** Monovalent affinities and multivalent apparent affinities of ETV:IDS and IgG:IDS to hTfR were measured by surface plasmon resonance. Abbreviations: kon, association rate constant; koff, dissociation rate constant; KD, equilibrium dissociation constant. *The value reported for the multivalent interaction between hTfR and IgG:IDS represents an apparent affinity. The complex binding kinetics for the multivalent IgG:IDS prevented binding kinetics from being fit (n/a). Graph displays mean values across all experimental replicates ± SEM.

The activities of ETV:IDS and IgG:IDS were assessed in MPS II patient-derived fibroblasts using an established ^35^S pulse-chase assay, in which ^35^S is integrated into newly-synthesized GAGs (Lu et al., 2010; Ullman et al., 2020). These fibroblast lines showed significant accumulation of heparan and dermatan sulfate and led to an ~2.5-fold accumulation of ^35^S signal (Ullman et al., 2020). Both ETV:IDS and IgG:IDS were highly active in reducing accumulation of 35S-labelled substrates with cellular IC_50_ values of 7.5 and 20 pM respectively (Fig. 1C). The higher activity of ETV:IDS in a cellular context was surprising given that IgG:IDS was 1.5-fold more active with respect to *in vitro* specific activity; however, cellular activity represents a complex integration of cell binding and uptake (through TfR and mannose-6-phosphate receptor (M6PR) interactions), delivery to the lysosome, and enzymatic activity within the lysosomal environment. We therefore further characterized the interactions with TfR and M6PR to better understand what might underlie differences in our *in vitro* and cellular results.

IDS functionality (including biodistribution) is strongly influenced by the incorporation of mannose-6-phosphate (M6P) on the terminal branches of N-linked glycosylations and its ability to bind to M6PR. To confirm that the different architectures retained strong M6PR binding, the affinities of ETV:IDS and IgG:IDS were determined using an enzyme-linked immunosorbent assay (ELISA). ETV:IDS bound recombinant M6PR with a median effective concentration (EC_50_) of 140 pM while IgG:IDS bound with an EC_50_ of 75 pM (Fig. 1D). The ~2-fold difference in M6PR affinity may reflect multivalent receptor interactions of the two IDS enzymes in IgG:IDS (compared to one in ETV:IDS). As M6PR provides an essential trafficking pathway for targeting enzymes from the extracellular space to the lysosome (El-Shewy and Luttrell, 2009; Urayama, 2013), the strong affinities of both ETV:IDS and IgG:IDS for M6PR suggest the molecules are functionally similar with respect to M6PR binding (ELISA, Fig. 1D) and therefore their M6PR-related trafficking is expected to be similar.

Interaction of both architectures with TfR is a critical attribute aimed at enabling brain uptake. Binding to TfR for ETV:IDS and IgG:IDS was assessed using two methods: monovalent affinities were determined by measuring binding to the soluble apical domain of human TfR (TfR^apical^), while apparent affinities arising from potential avid interactions with homodimeric TfR on the cell surface were approximated by measuring binding to full-length TfR receptor immobilized at a high density. ETV:IDS displayed affinities between 100 – 200 nM regardless of assay format (Fig. 1E), consistent with its ability to engage a single TfR. IgG:IDS can bind TfR bivalently and was strongly influenced by receptor density, having an affinity of 2.6 nM to TfR^apical^ and an apparent affinity of <100 pM to the full-length receptor (Fig. 1E). These values highlight the TfR affinity differences between the platforms and are in close agreement with previously reported results for ETV:IDS (Ullman et al., 2020) and for an anti-TfR antibody IDS fusion protein (Sonoda et al., 2018). Overall, IgG:IDS exhibited a monovalent affinity that was ~80-fold higher for TfR than ETV:IDS and the potential to bind TfR bivalently, leading to an apparent affinity ~1000-fold greater than ETV:IDS.

The *in vitro* attributes quantified here between ETV:IDS and IgG:IDS demonstrate that while there may be modest differences in activity and cellular potency, the most substantial difference between these platforms are in TfR affinity & valency. The approximately 2-fold reduction in cellular potency observed for IgG:IDS, despite its higher specific activity compared to ETV:IDS, is most likely due to the effect of architecture on cellular uptake and trafficking. We hypothesized that differences in TfR affinity and valency would result in significantly different CNS biodistribution and efficacy for ETV:IDS and IgG:IDS *in vivo*.

### ETV:IDS has enhanced peripheral exposure and improved brain uptake compared to IgG:IDS in TfR^mu/hu^KI mice

We next assessed the pharmacokinetic (PK) profiles of ETV:IDS and IgG:IDS in a chimeric mouse knock-in (KI) model expressing the human TfR^apical^ domain (TfR^mu/hu^ KI; (Kariolis et al., 2020)). Molecules with high-affinity to TfR such as IgG:IDS have historically been dosed at 1 – 3 mg/kg (Sonoda et al., 2018; Zhou et al., 2012), while low-affinity TfR binders have often been evaluated at much higher doses ranging from 20 – 50 mg/kg (Webster et al., 2017). These different dosing paradigms have led to the misperception that weaker TfR binding requires elevated doses to achieve brain exposures needed for a therapeutic response (Pardridge, 2015). Moreover, the reduced brain uptake often observed for high-affinity molecules at high doses has been suggested to be due to receptor saturation (Pardridge, 2015). It has been estimated that most LSDs likely only require a threshold activity of ~10% or less of normal residual enzyme activity to achieve full prevention of substrate storage and significant slowing of disease progression (Parenti et al., 2015). As long as enzyme formats achieve fairly uniform brain exposure that reliably extends beyond cerebral capillary endothelial cells, lower doses may therefore be sufficient to effectively reduce storage and disease progression.

To better understand the impact of dose on PK and biodistribution for the two platforms, TfR^mu/hu^ KI mice received an intravenous (IV) dose of either 1, 3, or 10 mg/kg ETV:IDS or IgG:IDS. Serum, liver, and brain concentrations of each molecule were determined to characterize biodistribution in key compartments (Ullman et al., 2020). Serum concentrations of ETV:IDS were elevated and more prolonged across dose levels compared to IgG:IDS (Fig. 2A) while IgG:IDS liver levels exceeded that of ETV:IDS (Fig. 2B), consistent with TfR mediated disposition from the circulation based on the high bivalent affinity of IgG:IDS. Importantly, ETV:IDS had 1.6- and 2.5-fold higher whole brain exposures compared to IgG:IDS at both 3 and 10 mg/kg, respectively (Fig. 2C and Table S1). Maximal brain concentrations for ETV:IDS was 1.7- and 2.7-fold higher compared to IgG:IDS at both 3 and 10 mg/kg, respectively at 8 – 24 hours post-dose (Fig. 2C and Table S1). In addition, IgG:IDS brain uptake appeared to plateau at 3mg/kg suggesting this architecture saturates its uptake mechanism at low concentrations and therefore does not display the dose-dependent brain uptake that is evident for ETV:IDS. Indeed, ETV:IDS continued to show non-saturable dose-dependent brain uptake up to 10 mg/kg (Fig. 2C). An effective delivery platform for IDS should ideally provide dose-dependent exposure capable of reaching therapeutic efficacy in both the periphery and brain. While the PK data for ETV:IDS displayed these attributes, IgG:IDS exhibited higher peripheral clearance and an upper limit on total brain exposure, thus illustrating how platform differences in TfR binding affinity and valency can impact *in vivo* biodistribution at the whole tissue level.

**Figure 2.**
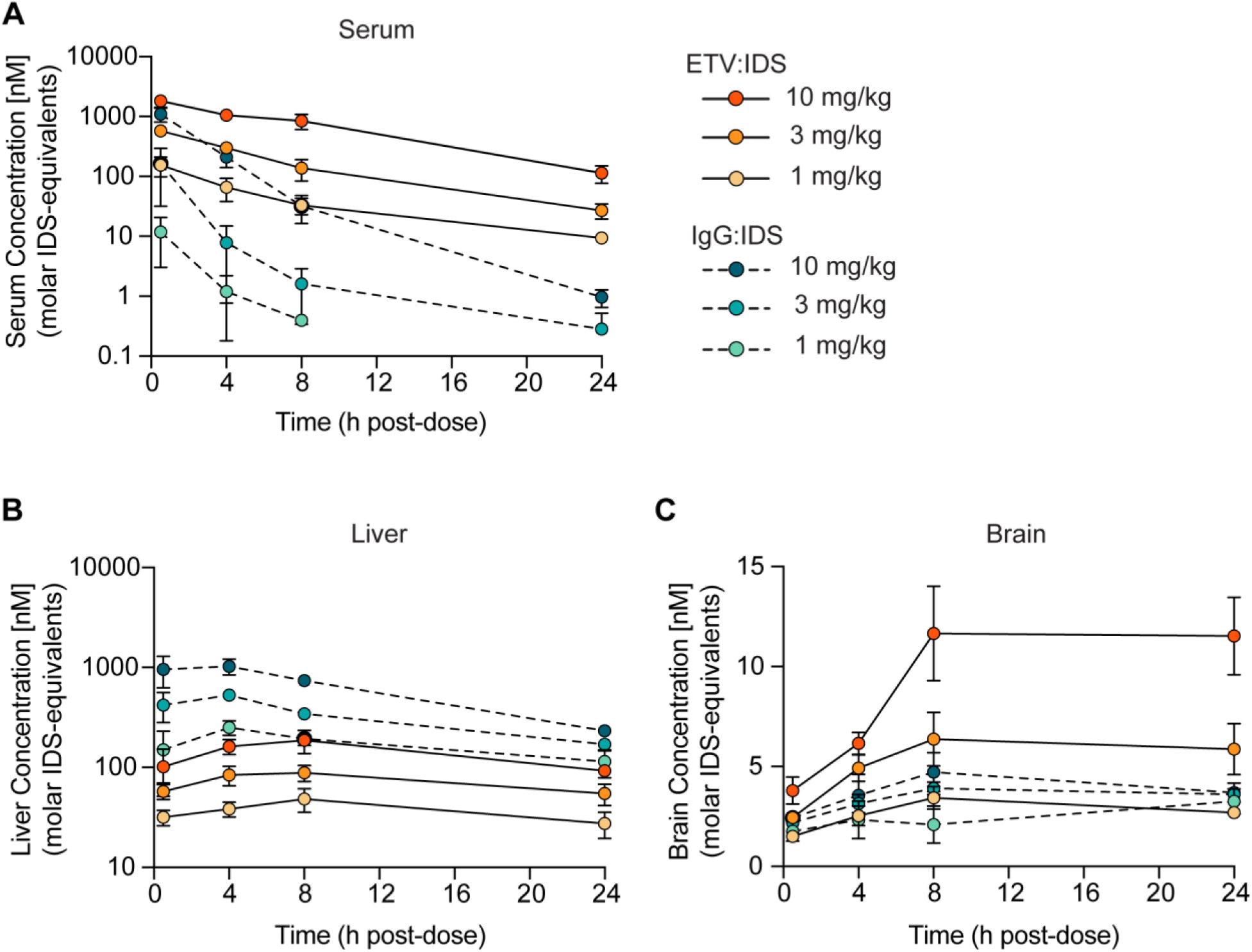
ETV:IDS has improved brain uptake compared to IgG:IDS in TfR^mu/hu^ KI mice. **(A)** Serum **(B)** liver and **(C)** brain concentrations of ETV:IDS or IgG:IDS from TfR^mu/hu^ KI mice were measured 0.5, 4, 8, and 24 hours after an intravenous dose of 1, 3 or 10 mg/kg and determined by IDS/IDS immunoassay; n = 3-5 per group. Graphs display mean ± SD.

### ETV:IDS distributes more effectively into the brain parenchyma than IgG:IDS in TfR^mu/hu^KI mice

A limitation in assessing biodistribution to the brain using whole tissue lysates (Fig. 2C) is that this approach does not differentiate between accumulation in the vasculature versus the parenchyma, making it difficult to distinguish actual transcytosis at the BBB from entrapment in capillary endothelial cells. Since it is well established that TfR-targeted proteins and nanoparticles may become trapped within brain capillary endothelial cells under some conditions (Bien-Ly et al., 2014; Johnsen et al., 2017; Manich et al., 2013; Moos and Morgan, 2001; Niewoehner et al., 2014; Paris-Robidas et al., 2011; Webster et al., 2017; Yu et al., 2011), we further assessed the brain biodistribution of ETV:IDS and IgG:IDS using brain capillary depletion. The capillary depletion method isolates brain vasculature from whole brain homogenate, allowing for a concentration determination in separate vascular and parenchymal fractions (Triguero et al., 1990). We utilized this method to evaluate the distribution of ETV:IDS and IgG:IDS into the brain 0.5, 4, and 24 hours after an IV dose in TfR^mu/hu^ KI mice (10 mg/kg). Notably, IDS concentrations in the brain vascular fraction for IgG:IDS increased at each timepoint whereas a minimal change for ETV:IDS was observed over time (Fig. 3A). In contrast, IDS levels in the parenchymal fraction steadily increased with time for ETV:IDS while IgG:IDS levels remained low over the entire 24 hours, resulting in 3.5- and 4-fold higher concentrations obtained with ETV:IDS compared to IgG:IDS at 4 and 24 hours post-dose (Fig. 3B). Expressing the data as a ratio of the parenchymal-to-vascular concentrations provided a further measure of distribution into the parenchyma, taking into account the IDS level in each fraction simultaneously (Fig. 3C). This ratio increased significantly over time for ETV:IDS with the 4 and 24 hour time points 26% and 165% higher, respectively, than the initial 0.5 hour value, while the IgG:IDS ratio was significantly lower than the ETV:IDS ratio at 0.5 hours and failed to increase over time (Fig. 3C). The low brain parenchymal fraction measured over 24 hours for IgG:IDS is consistent with other reports using a similar fusion protein (Sonoda et al., 2018). The data suggest that ETV:IDS effectively crossed the BBB with subsequent distribution into the brain parenchyma, while IgG:IDS was primarily trapped within the brain vasculature.

**Figure 3.**
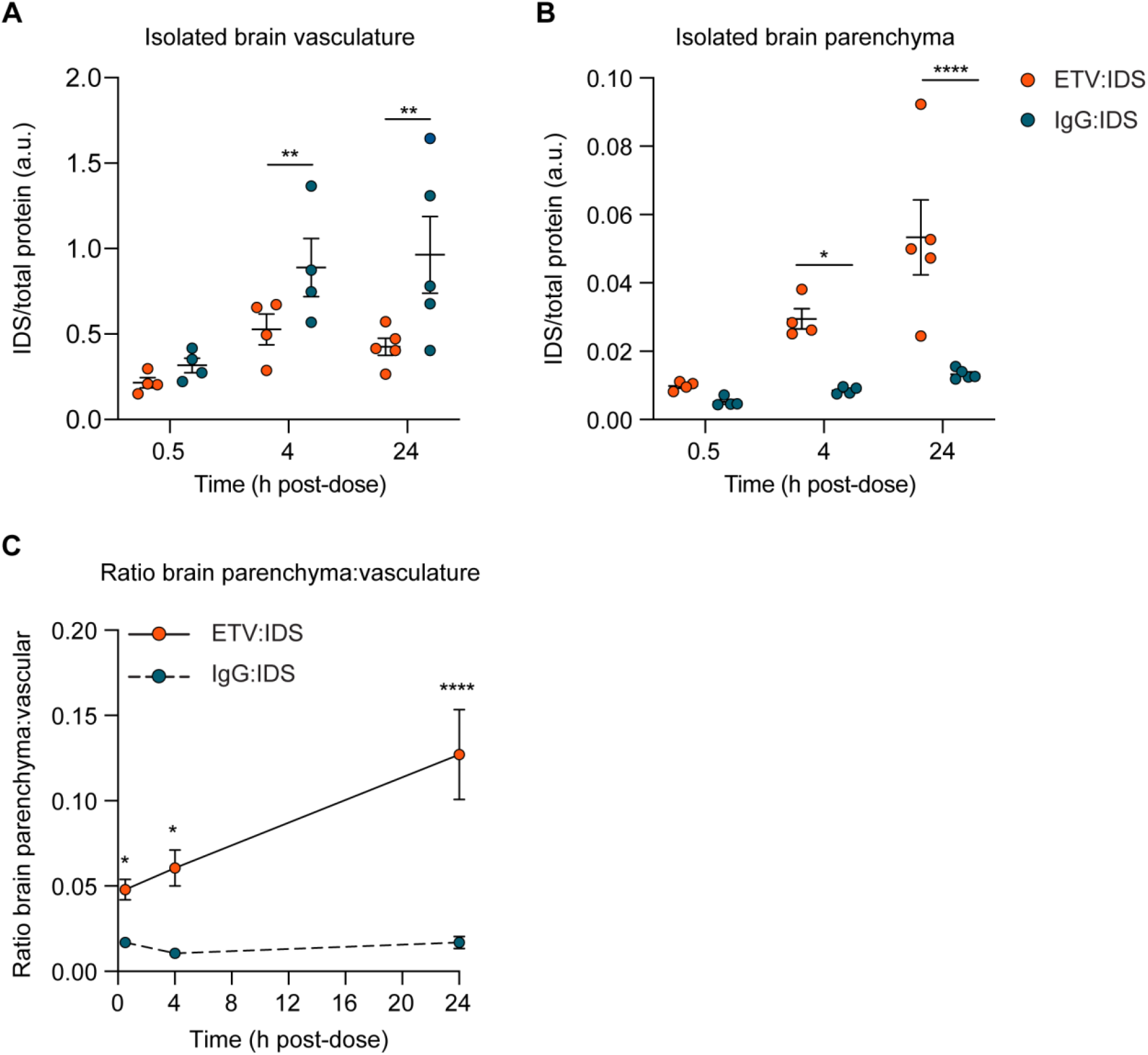
Capillary depletion method demonstrates accumulation of ETV:IDS in the brain parenchyma and IgG:IDS in the brain vasculature. The IDS concentrations of ETV:IDS or IgG:IDS in the isolated brain vasculature **(A)** or brain parenchymal fraction **(B)** from TfR^mu/hu^ KI mice measured at 0.5, 4, and 24 hours after an intravenous dose of 10 mg/kg. **(C)** Ratio of brain parenchyma to brain vasculature concentrations; n = 4-5 per group. Graphs display mean ± SEM and p values: two-way ANOVA with Sidak’s multiple comparison test; * p ≤ 0.05, ** p ≤ 0.01, and **** p ≤ 0.0001.

The precise molecular mechanisms underlying TfR-mediated transcytosis remain to be fully elucidated (Villasenor et al., 2019). Bivalent receptor interactions appear to drive TfR cross-linking and clustering at the brain endothelial cell luminal membrane, increasing uptake and lysosomal degradation while decreasing recycling, due partly to differential sorting within the endocytic pathway (Niewoehner et al., 2014; Villasenor et al., 2017; Weflen et al., 2013). High-affinity interactions with TfR, regardless of valency, also appear to lead to increased lysosomal trafficking of biotherapeutics in brain endothelial cells (Bien-Ly et al., 2014; Haqqani et al., 2018; Yu et al., 2011). Our capillary depletion data are consistent with these mechanisms. We show with ETV:IDS that decreasing affinity and restricting TfR binding to a monovalent interaction resulted in significantly higher distribution into the brain parenchyma, compared to IgG:IDS which distributed primarily within the brain vasculature.

### Imaging demonstrates that ETV:IDS enables greater parenchymal distribution, neuronal uptake and trafficking to parenchymal lysosomes than IgG:IDS in TfR^mu/hu^KI mice

Quantitative imaging methods and super resolution confocal microscopy (SRCM) were used to confirm the capillary depletion results and to better understand the brain distribution of ETV:IDS and IgG:IDS at the cellular and subcellular level. Imaging was performed on sagittal brain sections from TfR^mu/hu^ KI mice at 0.5, 4, and 24 hours after an IV dose (10 mg/kg). Brain sections were immunostained for human IgG (huIgG) and imaging was performed on multiple different brain regions (Fig. 4A and Fig. S1). Qualitative assessment of huIgG staining in the cortex of mice after administration of IgG:IDS exhibited a predominantly vascular pattern at all time points (Fig. 4B). Conversely, following ETV:IDS administration, huIgG staining in the cortex of mice was predominantly vascular at 0.5 hours but transitioned to a broad, diffuse parenchymal pattern with prominent cellular internalization by 24 hours (Fig. 4B). Quantitative analysis of extravascular huIgG staining in the cortex confirmed ETV:IDS signal accumulation in the brain parenchyma over time with the 4 and 24 hour time points 30% and 54% higher, respectively, than the initial 0.5 hour value (Fig. 4C). This contrasted with a relatively low and constant extravascular huIgG signal for IgG:IDS (Fig. 4C), in general agreement with capillary depletion results. A similar staining pattern and quantification was also observed for both molecules in all other regions assessed, including the hippocampus, hindbrain, and cerebellum (Fig. S1).

**Figure 4.**
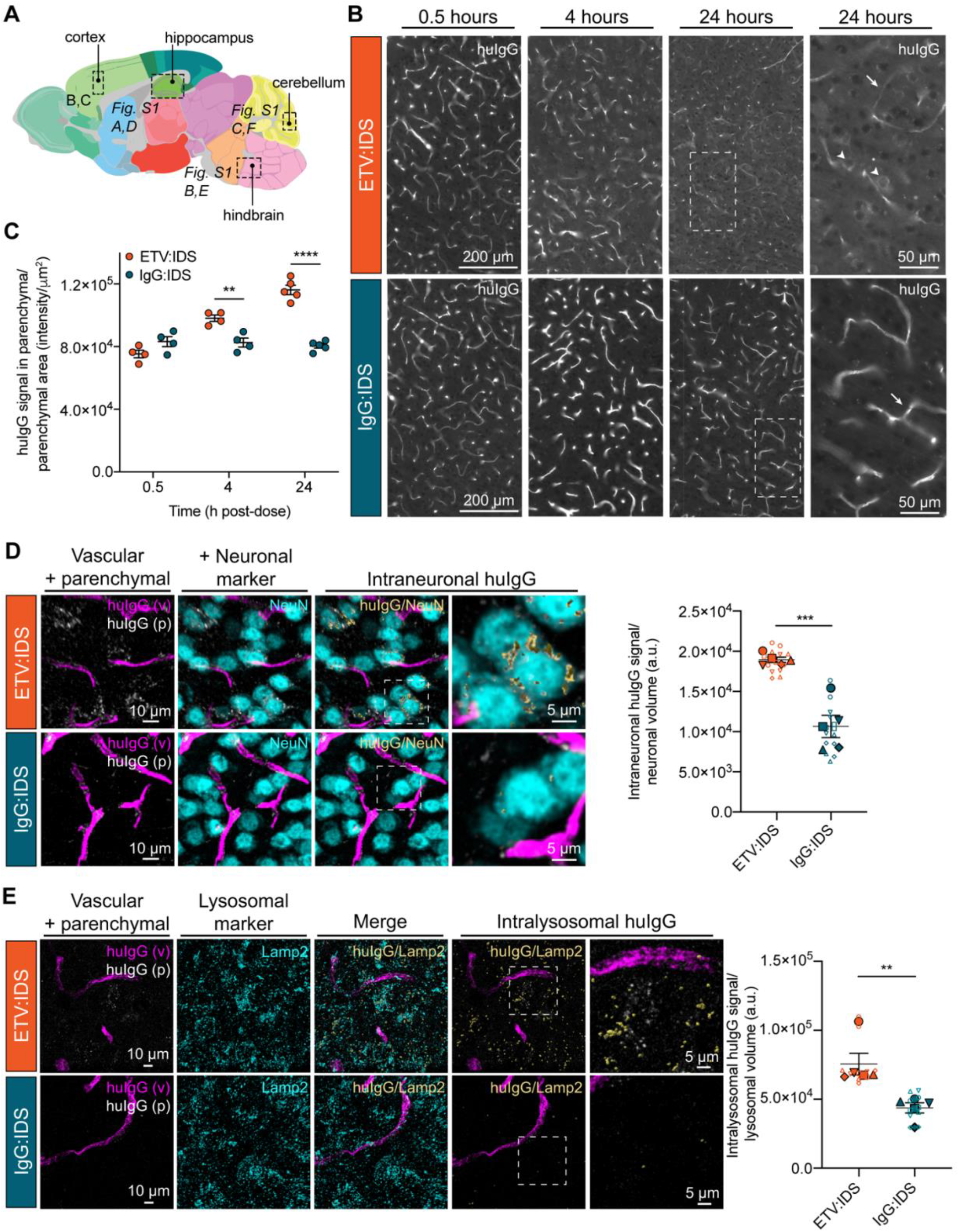
Immunohistochemical localization of ETV:IDS demonstrates enhanced distribution into the brain parenchyma compared to IgG:IDS. The distribution of ETV:IDS and IgG:IDS was assessed using sagittal brain sections from TfR^mu/hu^ KI mice at 0.5, 4, and 24 hours after an intravenous dose of 10 mg/kg. **(A)** Schematic indicating approximate location of sagittal brain regions of interest (ROI) for images and quantification in B-C (cortex) and Figure S1 (hippocampus, hindbrain, and cerebellum) (adapted from the Allen Adult Mouse Brain Atlas; original image credit: Allen Institute; E.S. Lein et al. Genome-wide atlas of gene expression in the adult mouse brain. *Nature*, 445:168-176, 2007). **(B)** Sections were immunostained with antibodies against huIgG and imaged using a wide field fluorescence slide scanner. Dashed boxes indicate regions shown at higher magnification displayed in the far-right panel. Arrows indicate huIgG staining localized to vascular profiles while arrowheads indicate cellular internalization of huIgG staining. **(C)** Quantification of huIgG staining in the parenchyma of the cortex was calculated based on the total sum intensity of all parenchymal staining in the ROI divided by the total parenchymal area in the ROI. A custom macro script was used to identify blood vessels present in the tissue and masked out of subsequent image analyses; n = 4-5 per group. Graphs display mean ± SEM and p values: two-way ANOVA with Sidak’s multiple comparison test; ** p ≤ 0.01 and **** p ≤ 0.0001. Sagittal brain sections from TfR^mu/hu^ KI mice at 24 hours after an intravenous dose of 10 mg/kg were immunostained with antibodies against (**D**) huIgG and the neuronal marker NeuN or (**E**) huIgG and the endo-lysosomal marker LAMP2. Confocal Z-stacks were acquired using a super resolution scanning confocal microscope. For the analysis, huIgG positive signal was segmented into vascular (magenta) and parenchymal (greyscale) components. (**D**) Intraneuronal huIgG (yellow, shown with surface rendering) and (**E**) intralysosomal huIgG (yellow, shown with surface rendering) was further segmented using either NeuN or LAMP2 (cyan) as a mask. **(E)** The merged panel shows both LAMP2 signal non-colocalized (cyan) and colocalized (yellow, shown with surface rendering) with huIgG signal. For better visualization, the subsequent intralysosomal panels show only the LAMP2 signal that colocalized with huIgG (yellow). The (**D**) intraneuronal and (**E**) intralysosomal huIgG signal was quantified and normalized to the total neuronal volume or total lysosomal volume, respectively. Graphs display superimposed summary statistics from 5 animals (solid shapes) consisting of 2-3 different image volumes from each animal (open shapes). Each animal is coded by different shapes. The 5 means were then used to calculate the mean ± SEM and p values: unpaired t test analysis; ** p ≤ 0.01 and *** p ≤ 0.001.

SRCM was used to investigate the subcellular localization of huIgG staining in brain sections from the cortex of TfR^mu/hu^ KI mice following an IV dose (10 mg/kg) of ETV:IDS or IgG:IDS at 24 hours post-dose, when staining differences were most pronounced. Imaging at subcellular resolution and subsequent segmentation of huIgG positive signal into vascular and parenchymal components revealed a more prominent, diffuse parenchymal staining pattern for ETV:IDS compared to IgG:IDS (Fig. 4D, E; left-most panels). Treatment with ETV:IDS also resulted in significantly greater huIgG signal in NeuN-positive cortical neurons compared to IgG:IDS (Fig. 4D); our use of SRCM demonstrating robust uptake of ETV:IDS into neurons is consistent with prior results reporting ETV:IDS uptake and effects in neurons using flow cytometry-based methods (Ullman et al., 2020). As lysosomal storage disorders lead to progressive substrate accumulation and perturbed lysosomal function, we also compared parenchymal cell internalization and trafficking to lysosomes for ETV:IDS and IgG:IDS. Consistent with increased distribution to cortical neurons, treatment with ETV:IDS resulted in significantly greater cortical huIgG signal in LAMP2-positive endo-lysosomes as compared to IgG:IDS. (Fig. 4E).

Taken together, our imaging results demonstrated that ETV:IDS was superior to IgG:IDS in achieving brain exposure beyond the vasculature, with broad distribution across brain regions, internalization into neurons, and lysosomal trafficking in brain cells. In contrast, IgG:IDS was predominantly within the brain vasculature, resulting in limited uptake into neurons and poor lysosomal trafficking in brain cells, thus showing that architecture determines biodistribution.

### ETV:IDS dose-dependently reduces brain GAG levels compared to IgG:IDS in Ids KO; TfR^mu/hu^ KI mice

We next investigated whether the higher whole brain exposures with ETV:IDS compared to IgG:IDS in TfR^mu/hu^ KI mice translated to more effective CNS GAG reduction in a mouse model of MPS II (*Ids* KO;TfR^mu/hu^ KI; (Ullman et al., 2020)). Mice received an IV dose of ETV:IDS or IgG:IDS at 1, 3, or 10 mg/kg, and GAG levels in liver, CSF, and brain were assessed after seven days. Both ETV:IDS and IgG:IDS were highly effective peripherally, reducing liver GAGs to TfR^mu/hu^ KI levels (Fig. 5A). In the CSF, ETV:IDS dose-dependently lowered GAGs by 68-80% compared to the vehicle treatment group, while IgG:IDS decreased CSF GAGs by 32-61% over the same dose range (Fig. 5B and Table S2). In brain tissue lysate, ETV:IDS significantly lowered GAG concentrations over the entire dose range (49-76% reduction), while IgG:IDS decreased GAGs to a substantially lesser extent (25-43%) (Fig. 5C and Table S2).

**Figure 5.**
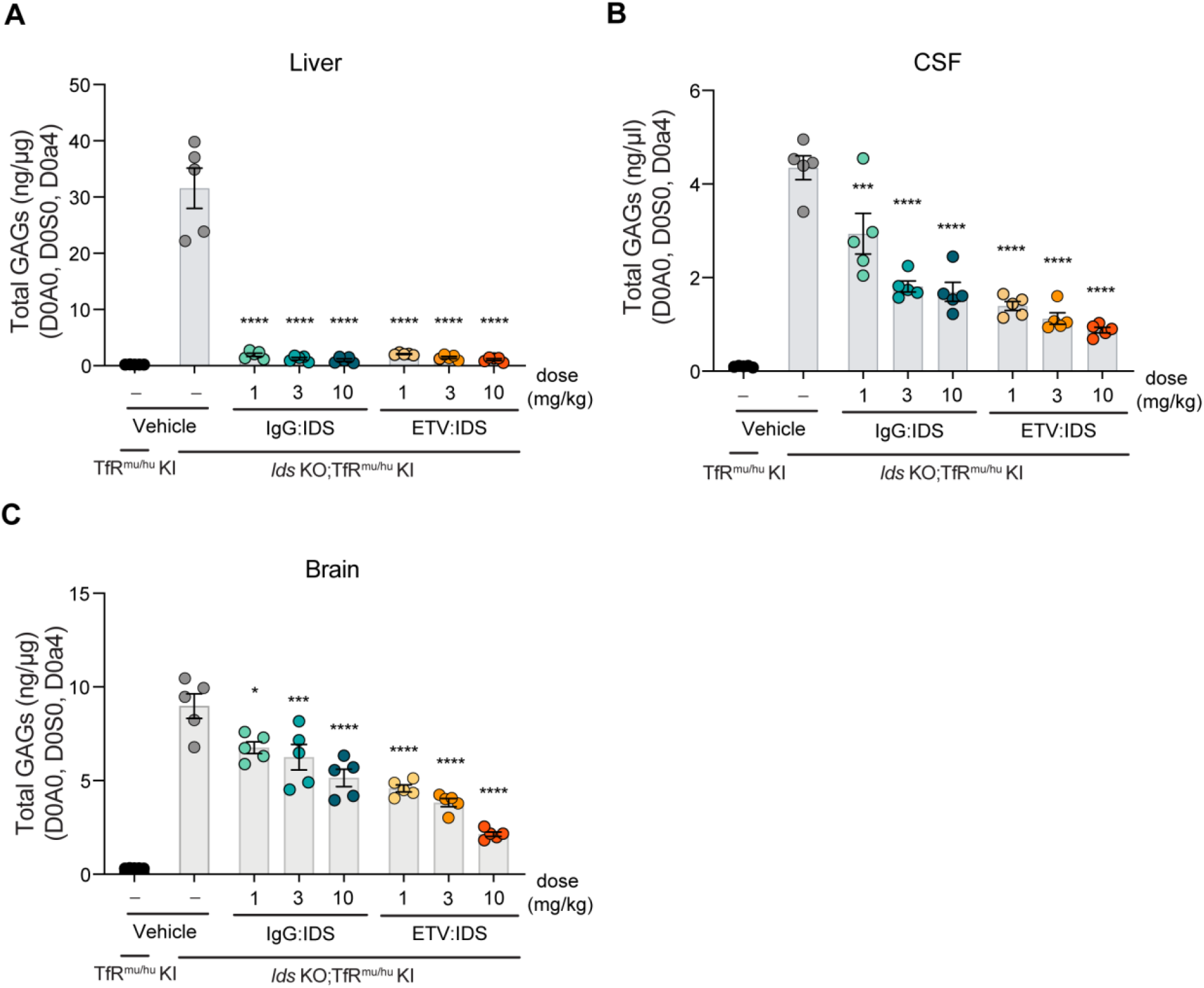
ETV:IDS is more effective than IgG:IDS at reducing brain and CSF GAGs in *Ids* KO;TfR^mu/hu^ KI mice. GAG levels were evaluated in the **(A)** liver, **(B)** CSF and **(C)** brain of *Ids* KO;TfR^mu/hu^ KI mice 7 days following treatment with ETV:IDS or IgG:IDS after an intravenous dose of 1, 3, or 10 mg/kg and compared to vehicle treatment and non-diseased TfR^mu/hu^ KI mice; n = 5 per group. Graphs display mean ± SEM and p values: one-way ANOVA with Tukey’s multiple comparison test; * p ≤ 0.05, *** p ≤ 0.001, and **** p ≤ 0.0001.

Taken together, this head-to-head comparison of two clinically relevant IDS fusion protein architectures, provides new evidence indicating how differences in TfR engagement can lead to significant and divergent effects on brain distribution and, ultimately, pharmacodynamics *in vivo.* Significant brain GAG reduction over the 1-10 mg/kg dose range for ETV:IDS was consistent with its superior biodistribution to the brain in TfR^mu/hu^ KI mice. Unlike IgG:IDS, ETV:IDS appeared to undergo pronounced transcytosis across brain endothelial cells to more effectively reach neurons and lysosomal compartments within the brain parenchyma, building upon recent evidence using other methods (Ullman et al., 2020). For IgG:IDS, the lack of dose-dependent brain uptake and GAG reduction in mouse models is consistent with poor trafficking across the BBB and inferior biodistribution to brain cells and limits the feasibility of exploring higher dose levels with an IgG:IDS architecture. The brain GAG reduction differences observed between ETV:IDS and IgG:IDS also agree well with our capillary depletion and imaging data, suggesting TfR engagement-dependent effects are most likely responsible for their differing efficacy. The size difference between ETV:IDS (~110 kDa) and IgG:IDS (~265 kDa) also may have subtly affected their relative biodistribution in brain extracellular space within and beyond the vascular basal lamina, although interstitial diffusion outward from capillaries is generally not expected to be highly limiting for molecules of this size (Wolak et al., 2015; Wolak and Thorne, 2013).

Our results also suggest CSF biomarker changes such as GAG reduction may accurately reflect pharmacodynamic effects in the brain only for IDS platforms where both preclinical distribution to brain parenchymal cells and corresponding effects on brain GAG reduction can be convincingly demonstrated, as reported here for ETV:IDS. Ultimately, further definitive insights into the relative advantages of diverse TfR-targeting brain delivery platforms may only be provided by the demonstration of efficacy in human patients. A Phase I/II clinical trial studying the ETV:IDS approach (https://clinicaltrials.gov/ct2/show/NCT04251026) and a Phase II/III clinical trial with an approach utilizing an IgG:IDS-like architecture (https://clinicaltrials.gov/ct2/show/NCT03568175) are currently in progress with clinical readouts expected in the near future. From the non-clinical data described here, understanding the impact of binding mode between the biotherapeutic and the receptor may ultimately prove to be a key consideration in successful clinical translation.

## Materials and Methods

### Cloning and architecture of ETV:IDS and IgG:IDS

All material used in this study was produced for research purposes. ETV:IDS was expressed as a knob-in-hole recombinant fusion protein consisting of IDS fused at the C-terminus (via a peptide linker) to a single copy of human Fc engineered to bind hTfR (Kariolis et al., 2020). IgG:IDS was expressed as a recombinant fusion protein consisting of an anti-TfR mAb fused at the C-terminus (via a peptide linker) of each heavy chain to IDS (Patent: CAS#2140211-48-7). Antibody constant regions used were human IgG1. Mutations mitigating effector function (L234A, L235A) were included on ETV:IDS (Kariolis et al., 2020) whereas effector function was maintained on IgG:IDS as described (L234, L235) (Sonoda et al., 2018).

### Protein expression

The IgG:IDS construct was expressed via transient transfection of ExpiCHO cells (Thermo Fisher Scientific) according to manufacturer’s instructions. Cultures were co-transfected with plasmids encoding IgG:IDS (heavy chain and light chains) and varying ratios of SUMF1 cDNAs. ETV:IDS was purified from a stable CHO clone.

### Protein purification

ETV:IDS and IgG:IDS were purified to homogeneity from serum-free CHO cultures by a series of chromatographic steps. IgG:IDS was affinity purified using Protein A followed by anion-exchange chromatography. Final fractions with a high degree of purity (as assessed by analytical size-exclusion chromatography (SEC) and/or microcapillary electrophoresis) were pooled, concentrated and dialyzed into 20mM sodium phosphate pH6.5 and 130mM NaCl. ETV:IDS was purified by a three-step chromatographic approach: Protein A affinity followed by anion exchange and finally hydrophobic interaction chromatography. Preparations were stored at 4°C or −80°C prior to use and routinely analyzed by SEC, specific activity and for endotoxin content.

### Biochemical Characterization

#### K_D_ determination using Surface Plasmon Resonance

The affinity of ETV:IDS and IgG:IDS for hTfR were determined by surface plasmon resonance on a Biacore™ T200 instrument using two methods, similar to previously described (Kariolis et al., 2020). In Method 1, to evaluate monovalent TfR binding affinities ETV:IDS or IgG:IDS were captured on a Protein A-coated Biacore™ Series S CM5 sensor chip and serial dilutions of hTfR^apical^ were injected over the captured sample. In Method 2, for evaluation of apparent hTfR binding affinities of ETV:IDS and IgG:IDS, biotinylated full-length hTfR was immobilized on a streptavidin-coated CM5 sensor chip, followed by injection of serial dilutions of ETV:IDS or IgG:IDS. For both methods, the single-cycle kinetics mode was used for sample injection (association time, 90 s; dissociation time, 600 s), and binding affinities were calculated using Biacore T200 Evaluation Software v3.0.

### In vitro IDS activity assay

The specific activity of IDS containing constructs were measured with a two-step fluorometric enzymatic assay using an artificial substrate as previously described (Voznyi et al., 2001). In brief, a 4-Methylumbelliferone (leaving group) standard curve was fit by linear regression to calculate the amount of product and verified as less than 10% of total substrate cleavage. Specific activity calculated as pmol product/minute/mg of protein was determined.

### ELISA-based analysis of M6PR binding

ELISA plates were coated with M6PR Fc at 1 μg/mL in PBS overnight at 4°C. The following day, the plate was washed three times with wash buffer (PBS with 0.02% Tween-20) and blocking buffer (PBS with 0.02% Tween-20 and 5% BSA) was added to each well. Blocking was carried out for 1 hour at room temperature after which the plate was washed three times, and ETV:IDS or IgG:IDS were added to the first column of the plate at a concentration of 25 nM. A 3-fold serial dilution was performed across the plate. Primary incubation of the binding reactions was done for 1 hour at room temperature. After binding, the plate was washed three times, and binding was detected using biotinylated anti-IDS antibody diluted to 0.0625 μg/mL in sample buffer. The plate was incubated with detection antibody for 1 hour at room temperature and then washed three times. Streptavidin-HRP, diluted 1:50,000 in sample buffer, was then added to each well. The plate was incubated for 30 minutes at room temperature and then washed three times. The ELISA was developed using TMB reagent. The ELISA plate was read on a HighRes BioTek Synergy plate reader, where the absorbance at 450nm was recorded.

### S^35^-sulfate accumulation assay to assess cellular potency

Cellular potency of IgG:IDS and ETV:IDS was carried out as described previously (Ullman et al., 2020). MPS II patient (GM01928, GM12366, GM13203) primary fibroblasts, were obtained from Coriell. The cellular S^35^-accumulation assay was performed using a method modified from Lu and co-workers (Lu et al., 2010). Briefly, fibroblasts were plated at 25,000 cells/well in 96-well plates and grown in DMEM high glucose (Gibco) with 10% FBS (Sigma). After 3 days of culture, media was replaced with low-sulfate F12 medium (Gibco) supplemented with 10% dialyzed fetal bovine serum and 40 mCi/mL [S35] sodium sulfate (PerkinElmer) for 96 hours. Following [S35] sodium sulfate incubation, cells were treated with ETV:IDS or IgG:IDS. After 24 hours of incubation, media was aspirated, cells were washed with cold PBS, and lysed with 0.01 N NaOH. Incorporated S35 was measured by scintillation counting (Microbeta Trilux). EC_50_ curves were generated using Prism software using a log(agonist) vs. response, variable slope (four parameter) fit.

### Animal care

All procedures in animals were performed in adherence to ethical regulations and protocols approved by Denali Therapeutics Institutional Animal Care and Use Committee. Mice were housed under a 12-hour light/dark cycle and had access to water and standard rodent diet (LabDiet® #25502, Irradiated) *ad libitum*.

### Mouse strains

A previously described *Ids* KO mouse model on a B6N background were obtained from Jackson Laboratories, JAX strain 024744 (Muenzer et al., 2002). The TfR^mu/hu^ KI mouse line harboring the human TfR apical domain knocked into the mouse receptor was developed by generating a knock-in (into C57Bl6 mice) of the human apical TfR mouse line via pronuclear microinjection into single cell embryos, followed by embryo transfer to pseudo pregnant females using CRISPR/Cas9 technology. The donor DNA comprised the human TfR apical domain coding sequence that has been codon optimized for expression in mouse. The resulting chimeric TfR was expressed *in vivo* under the control of the endogenous promoter. A founder male from the progeny of the female that received the embryos was bred to wild-type females to generate F1 heterozygous mice. Homozygous mice were subsequently generated from breeding of F1 generation heterozygous mice (Kariolis et al., 2020). TfR^mu/hu^ KI male mice were bred to female *Ids* heterozygous mice to generate *Ids* KO; TfR^mu/hu^ KI mice (Ullman et al., 2020). All mice used in this study were males.

### Biodistribution and pharmacokinetics of ETV:IDS and IgG:IDS

ETV:IDS or IgG:IDS were administered intravenously via the tail vein to 2-3 month old TfR^mu/hu^ KI mice (n = 3-5 per group) at doses of 1, 3, or 10 mg/kg body weight and animals were sacrificed at 0.5, 4, 8, and 24 hours post-dose. For terminal sample collection, animals were deeply anesthetized via intraperitoneal (i.p.) injection of 2.5% Avertin. Blood was collected via cardiac puncture for serum collection and allowed to clot at room temperature for at least 30 minutes. Tubes were then centrifuged at 12,700 rpm for 7 minutes at 4°C. Serum was transferred to a fresh tube and flash-frozen on dry ice. Animals were transcardially perfused with ice-cold PBS using a peristaltic pump (Gilson Inc. Minipuls Evolution) and the liver and brain were dissected. Liver and brain tissue (50 mg) were flash-frozen on dry ice and processed for an IDS/IDS ELISA as described below. Brain tissue from the 10 mg/kg groups were processed for capillary depletion and immunohistochemistry, as described below.

### Pharmacodynamics of ETV:IDS and IgG:IDS

ETV:IDS or IgG:IDS were administered intravenously via the tail vein to 2-3 month old TfR^mu/hu^ KI (n = 5 per group) and *Ids* KO;TfR^mu/hu^ mice (n = 5 per group) at doses of 0, 1, 3, or 10 mg/kg body weight and animals were sacrificed 7 days post-dose. For terminal sample collection, animals were deeply anesthetized via i.p. injection of 2.5% Avertin. For CSF collection, a sagittal incision was made at the back of the animal’s skull, subcutaneous tissue and muscle was separated to expose the cisterna magna and a pre-pulled glass capillary tube was used to puncture the cisterna magna to collect CSF. CSF was transferred to a Low Protein LoBind Eppendorf tube and centrifuged at 12,700 rpm for 10 minutes at 4°C. CSF was transferred to a fresh tube and snap frozen on dry ice. Lack of blood contamination in mouse CSF was confirmed by measuring the absorbance of the samples at 420 nm. Blood, serum and tissues were obtained as described and flash frozen on dry ice.

### Tissue processing for PK analysis

Tissue (50 mg) was homogenized in 10x volume by tissue weight cold 1% NP40 lysis buffer (1mL 10% NP-40 Surfact-Amps detergent solution, 9mL 1xPBS, 1 tablet cOmplete protease inhibitor, 1 tablet PhosSTOP protease inhibitor) with a 3 mm stainless steel bead using the Qiagen TissueLyzer II for two rounds of 3 minutes at 27Hz. Homogenates were then incubated on ice for 20 minutes and spun at 14,000rpm for 20 minutes at 4°C. The resulting lysate was transferred to a single use aliquot and stored at −80°C.

### IDS:IDS ELISAs and PK Analysis

ETV:IDS and IgG:IDS were measured in serum, liver lysates, and brain lysates using an Iduronate-2-Sulfatase sandwich ELISA. A 384-well maxisorp plate (Thermo, Cat# 464718) was coated overnight with an anti-IDS antibody (R&D Systems, Cat# AF2449) and blocked with Casein-PBS Buffer (Thermo, Cat# 37528) the following day. Samples containing either ETV:IDS or IgG:IDS were added to the plate and incubated for one hour at room temperature. After a subsequent wash, a biotinylated anti-IDS antibody (R&D Systems, Cat # BAF2449) was added to bind the immobilized ETV:IDS and IgG:IDS. The IDS sandwich was then detected with a streptavidin-horseradish peroxidase conjugate (Jackson Immuno Research, Cat#016-030-084) followed by incubation with a 3,3’,5,5’tetra-methyl-benzidine (TMB) substrate (Thermo, Cat# 34028). The reaction was quenched with 4N hydrosulfuric acid (Life Technologies, Cat#SS04) and the plate was read at 450 nm absorbance wavelength on a plate spectrophotometer to determine the concentrations of analyte in the samples. Calibration standard curves were generated for ETV:IDS and IgG:IDS using a 5-Parameter Logistic Fit with an assay range of 0.00137 nM – 1 nM. Protein sequence-derived molecular weights were used to calculate molar concentrations. To compare ETV:IDS and IgG:IDS concentrations in terms of molar equivalents of IDS enzyme (Fig. 2), a correction factor was applied to IgG:IDS concentration data to account for the 2:1 ratio of IDS enzyme per mole of IgG:IDS (Fig. 1). Serum, liver, and brain area under the curve (AUC) exposures for ETV:IDS and IgG:IDS were calculated using non-compartmental analysis (NCA) in Dotmatics, version 4.8 (Bishop’s Stortford, UK). Semi-log and linear graphs and tabular results with standard deviations were prepared with Prism 8 (GraphPad, San Diego, CA).

### Capillary depletion

The meninges and choroid plexuses were removed from the brain pieces reserved for capillary depletion immediately following extraction to ensure that these blood-CSF barriers do not contribute to an overestimation of IDS concentration in the isolated brain parenchyma samples. The capillary depletion method was conducted as previously described (Kariolis et al., 2020). Briefly, fresh brain pieces were homogenized on ice by 10 strokes with of a Dounce homogenizer (smaller diameter pestle) in 3.5 mL HBSS then centrifuged for 10 min at 1,000 g. The pellet was resuspended in 2 mL of 17% dextran (MW 60,000; Sigma 31397) and centrifuged for 15 min at 4,122 g to separate the parenchymal cells from the vasculature. The top myelin and parenchymal cell layers were removed together and diluted with HBSS, then centrifuged for 15 min at 4,122 g to pellet the parenchymal cells. Both the vascular pellets and parenchymal cell pellets were resuspended in cold 1% NP40 in PBS with protease and phosphatase inhibitors (Sigma 04693159001 and 04906837001), agitated 30 s at 27 Hz with a Tissue Lyser II (Qiagen, 85300), then incubated 20 min on ice. The cell lysate was collected following centrifugation for 10 min at 12,700 g. The total protein concentration of each sample was measured using a BCA assay (Thermo 23225).

### Immunohistochemistry

Immediately following extraction, mouse brain tissue was fixed in 4% paraformaldehyde for 24 hours at 4°C then transferred to 30% sucrose in 1xPBS for at least 24 hours at 4°C. Sagittal tissue sections at 30 micron thickness were sectioned using a Leica Sliding Microtome. Free floating sections were collected in 2 mL Eppendorf Tubes filled with 1xPBS with 0.05% sodium azide and either directly mounted onto Fisherbrand Superfrost Plus microscope slides or used as free-floating sections throughout the IHC process. Sections were rinsed in 1xPBS for 2 rounds of 5 minutes then transferred to Sequenza Clips (for those sections directly mounted onto microscope slides) and rinsed in 1xPBS/0.05% Tween. Sections were then permeabilized in 0.5% Triton X-100 for 15 minutes and incubated in Blocking Solution (1%BSA/0.1% fish skin gelatin/0.5% Triton X-100/0.1% sodium azide in 1xPBS) for 2 hours at room temperature. Sections were then incubated in primary antibody (Jackson ImmunoResearch 109-605-003: Goat anti-huIgG conjugated to Alexa Fluor 647, 1:250, Abcam GL2A7: Rat anti-LAMP2, 1:500, Abcam EPR12763: Rabbit anti-NeuN, 1:500) prepared in Antibody Dilution Buffer (1%BSA/0.1% sodium azide in 1xPBS) overnight at 4°C. Sections were rinsed in 1xPBS/0.05% Tween for 3 rounds of 5 minutes followed by incubation in secondary antibody (Invitrogen: Goat anti-rat Alexa Fluor 555, 1:500 and Donkey anti-rabbit Alexa Fluor 488) prepared in Antibody Dilution Buffer for 2 hours at room temperature in the dark. Sections were then rinsed in 1xPBS/0.05% Tween for 3 rounds of 5 minutes and incubated in DAPI (Invitrogen Molecular Probes D1306: 1:10,000 from 5mg/mL stock) for 10 minutes. Sections were then rinsed in 1xPBS/0.05% Tween for 2 rounds of 5 minutes and either removed from the Sequenza Clips and quickly rinsed in 1xPBS or directly mounted onto Fisherbrand Superfrost Plus microscope slides for free floating sections. Sections were then cover slipped with Invitrogen Prolong Glass Antifade Mountant and cured overnight at room temperature.

### Image acquisition and quantification of CNS parenchymal antibody levels

Images of whole slide-mounted immunostained sagittal mouse brain sections were acquired using a wide field epifluorescence slide scanning microscope (Zeiss Axio Scan Z1; Carl Zeiss, Inc.), with a 40x/0.95 NA air objective and filter sets to specifically image Alexa 488, Alexa 555, and Alexa 647 labeled secondary antibodies. Exposure times were held constant for each channel across all fields / sections / slides imaged. Every field for each tissue section image was collected and post-stitched using Zeiss Zen Blue Edition software (v. 2.6). To quantify levels of huIgG present in the brain parenchyma of test subject animals, collected images were quantified using Zeiss Zen software. Specifically, a custom macro script was written to identify blood vessels present in the tissue, based on morphology and high levels of huIgG signal observed in the vessels of injected animals. Vessels were then masked out of subsequent image analyses, leaving only the surrounding parenchymal tissue to be quantified. Specific sub-regions of the brain sections were then selected for quantification (cortex, hippocampus, cerebellum, and hindbrain); positionally similar brain regions were selected across all quantified brain sections. Mean fluorescence intensities corresponding to detected test antibodies in selected brain parenchymal regions were then calculated based on the total sum intensity of all parenchymal (non-vessel) pixels in the selected regions divided by the total parenchymal area of the selected region.

### Super resolution confocal imaging and quantification of intracellular antibody levels

To quantify the intracellular localization of huIgG, sections were imaged using a scanning confocal microscope (Leica SP8; Leica Microsystems, Inc). operated in super resolution LIGHTNING mode, acquired with a 63x/1.4 NA oil objective at a pixel size of 50 nm and processed using the Adaptive processing algorithm. Confocal z-stacks of 25-30 μm were acquired for each channel using sequential scan settings from three independent cortical brain regions and from three animals in each treatment group. The huIgG signal within the intraneuronal compartment was masked using an intensity-based segmentation of NeuN positive pixels, and the resulting sum intensities were normalized to the total neuronal volume within a given three-dimensional image field. Intralysosomal huIgG was masked using an intensity-based segmentation of LAMP2 positive pixels and the huIgG sum intensities were quantified and normalized to the total lysosomal volume. In each case, the mean sum intensity was determined for each animal, and the 3 means were then used to calculate the mean ± SEM for each treatment.

### Tissue or fluid processing for GAG analysis

Tissue (50 mg) was homogenized in 750 μL water using the Qiagen TissueLyzer II for 3 minutes at 30Hz. Homogenate was transferred to a 96-well deep plate and sonicated using a 96-tip sonicator (Q Sonica) for 10×1 second pulses. Sonicated homogenates were spun at 2,500x*g* for 30 minutes at 4^°^C. The resulting lysate was transferred to a clean 96-well deep plate, and a BCA was performed to quantify total protein. 10 μg total protein lysate or 3 μL of CSF was used for subsequent HS/DS digestion. Digestion was carried out in a PCR plate in a total volume of 62 μL. Internal standard mix of HS and DS (20 ng total) were added to each sample and mixed with Heparinases I, II, III and Chondriotinase B in digestion buffer for 3 hours with shaking at 30^°^C. After the digest, EDTA was added to each sample and the mixture was boiled at 95^°^C for 10 minutes. The digested samples were spun at 3,364x*g* for 5 minutes and samples were transferred to a cellulose acetate filter plate (Millipore, MSUN03010) and spun at 3,364x*g* for 5 minutes. The resulting flow through was mixed with equal parts of acetonitrile in glass vials and analyzed by mass spectrometry as described below.

### Mass spectrometry analysis of GAGs

Quantification of GAG levels in fluids and tissues was performed by liquid chromatography (Shimadzu Nexera X2 system, Shimadzu Scientific Instrument, Columbia, MD, USA) coupled to electrospray mass spectrometry (Sciex 6500+ QTRAP, Sciex, Framingham, MA, USA). For each analysis, sample was injected on a ACQUITY UPLC BEH Amide 1.7 mm, 2.1×150 mm column (Waters Corporation, Milford, MA) using a flow rate of 0.6 mL/minute with a column temperature of 55°C. Mobile phases A and B consisted of water with 10 mM ammonium formate and 0.1% formic acid, and acetonitrile with 0.1% formic acid, respectively. An isocratic elution was performed with 80% B throughout the 8-minute run. Electrospray ionization was performed in the negative-ion mode applying the following settings: curtain gas at 20; collision gas was set at medium; ion spray voltage at −4500; temperature at 450°C; ion source Gas 1 at 50; ion source Gas 2 at 60. Data acquisition was performed using Analyst 1.6.3 or higher (Sciex) in multiple reaction monitoring mode (MRM), with dwell time (msec) for each species. Collision energy at −30; declustering potential at −80; entrance potential at −10; collision cell exit potential at −10. GAGs were detected as [M-H]^−^ using the following MRM transitions: D0A0 at m/z 378.1 > 87.0; D0S0 at m/z 416.1 > 138.0; D0a4 at m/z 458.1 > 300.0; D4UA-2S-GlcNCOEt-6S (HD009, Iduron Ltd, Manchester, UK) at m/z 472.0 (in source fragment ion) > 97.0 was used as internal standard.

Individual disaccharide species were identified based on their retention times and MRM transitions using commercially available reference standards (Iduron Ltd). GAGs were quantified by the peak area ratio of D0A0, D0S0, and D0a4 to the internal standard using Analyst 1.7.1 or MultiQuant 3.0.2 (Sciex). Reported GAG amounts were normalized to total protein levels as measured by a BCA assay (Pierce). Fold over TfR^mu/hu^ KI values were calculated as follows:

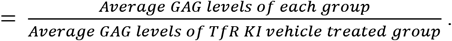

Percent reduction from vehicle treated *Ids* KO;TfR^mu/hu^ KI mice were calculated as follows:

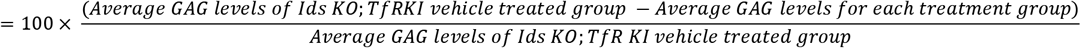

### Heparan sulfate (HS) and dermatan sulfate (DS) calibration curves

Pure standards for D0a4 (DS/CS), D0A0 (HS), and D0S0 (HS) were dissolved in acetonitrile:water 50/50 (v/v) to generate a 1 mg/mL stock. An 8-point dilution curve in PBS was generated ranging from 0.12 ng to 1000 ng. Subsequently, the internal standard D4UA-2S-GlcNCOEt-6S (20 ng) was added to each serial dilution. Samples were then boiled for 10 minutes at 95°C and then spun at 3,364xg to pellet any particulate matter. Supernatant was filtered using a 30kD MWCO cellulose acetate filter plate (Millipore, MSUN03010) by spinning at 3364xg for 5 minutes at room temperature. Resulting flow through was mixed with an equal part of acetonitrile in glass vials and run by mass spectrometry as described above.

## Author Contributions

**Conceptualization:** A.A., C.S.M., R.J.W., R.G.T., A.G.H., and M.S.K. **Methodology:** M.F. and B.T. **Software:** J.C.D. **Formal Analysis:** A.A, M.E.K.C., D.C., M.E.P., D. J. K., J.W., and R.G.T. **Investigation:** A.A., C.S.M., M.E.K.C., J.C.D., M.E.P., E.R.T., R.C., L.A.D., J.D., T.E., M.F., T.G., D.J.K., N.L., I.A.L., H.N.N., H.S., B.T., J.C.U., and M.S.K. **Writing - Original Draft:** A.A., C.S.M., M.E.K.C., M.E.P., R.G.T., A.G.H., and M.S.T. **Writing – Review and Editing:** M.S.D., K.R.H., and R.J.W. **Visualization:** A.A., C.S.M., M.E.K.C., M.E.P., R.G.T., A.G.H., and M.S.K. **Supervision:** A.A., C.S.M., M.S.D., D.D., K.G., K.R.H., J.W.L., P.E.S., M.D.T., J.M.H., K.S., L.S., R.J.W., R.G.T., A.G.H., and M.S.K.

