## Supplemental Tables and Figures for "Molecular architecture determines brain delivery of a transferrin-receptor targeted lysosomal enzyme"

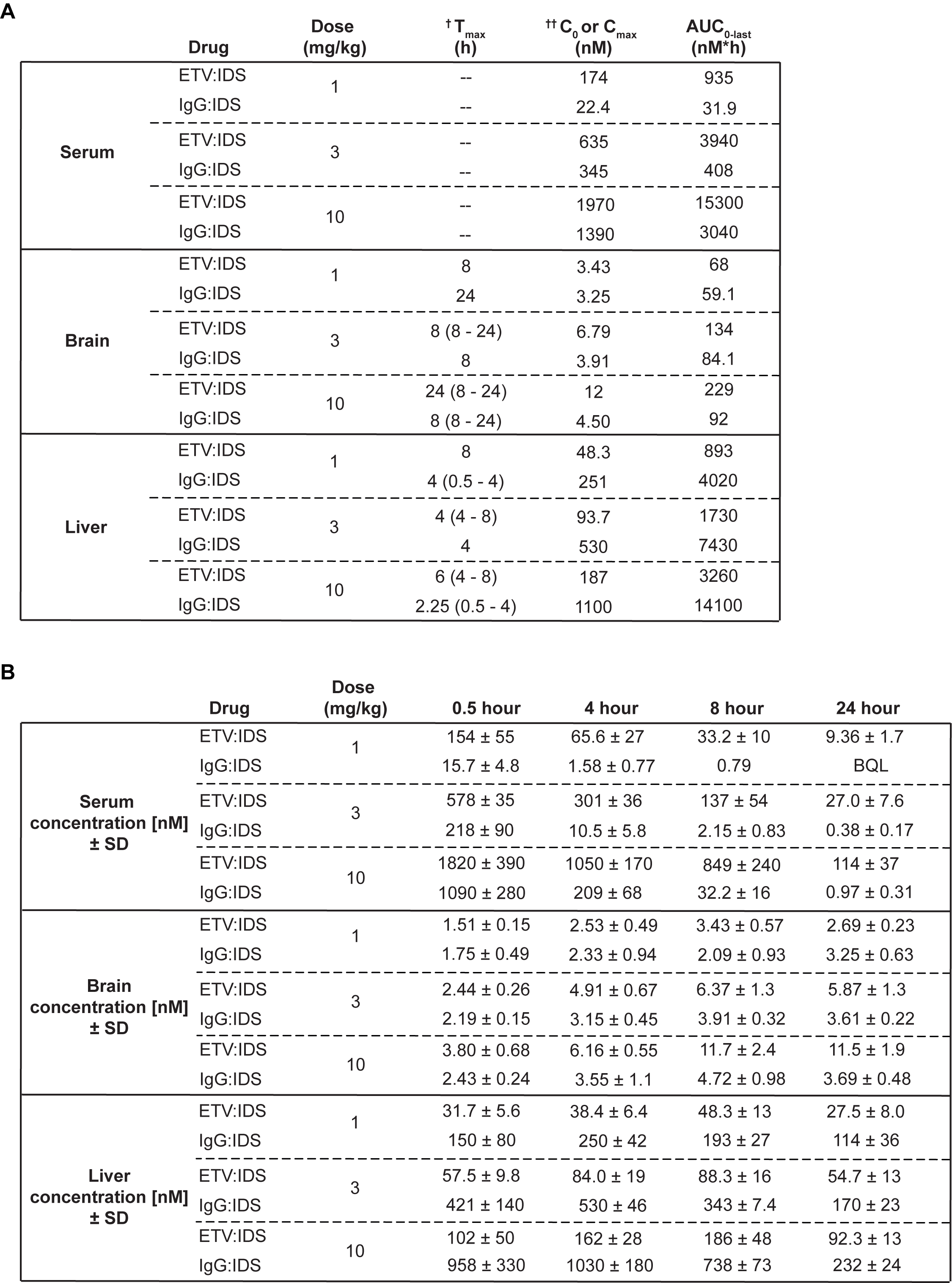

**Supplemental Table 1. Pharmacokinetic parameters. (A)** Serum, brain, and liver pharmacokinetic parameters at 1, 3, and 10 mg/kg of ETV:IDS or IgG:IDS. ^†^T_max_ is median (range) in brain and liver. ^††^C_0_ is indicated for serum whereas C_max_ is indicated for brain and liver. **(B)** Serum, brain and liver concentrations of ETV:IDS or IgG:IDS from TfR^mu/hu^ KI mice were measured 0.5, 4, 8, and 24 hours after an intravenous dose of 1, 3 or 10 mg/kg and determined by IDS/IDS immunoassay; n = 3-5 per group. Table displays mean values ± SD. BQL = below quantitation limit (Lower limit of quantitation = 0.00412 nM).

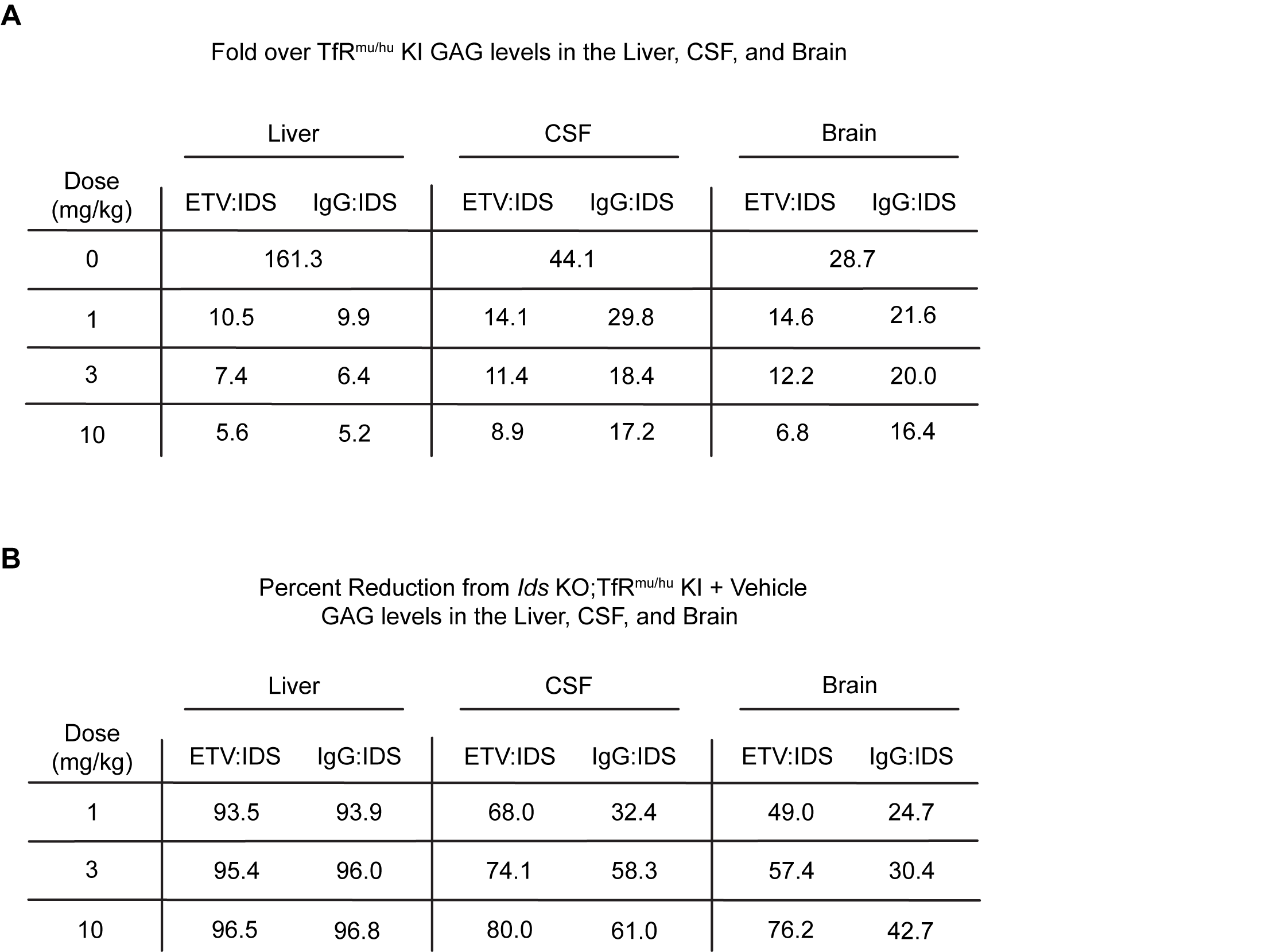

**Supplemental Table 2.** **ETV:IDS is more effective than IgG:IDS at reducing brain and CSF GAGs in *Ids* KO;TfR^mu/hu^ KI mice.** GAG levels were evaluated in the liver, CSF and brain of *Ids* KO;TfR^mu/hu^ KI mice 7 days following treatment with ETV:IDS or IgG:IDS after an intravenous dose of 1, 3, or 10 mg/kg and compared to vehicle treatment and non-diseased TfR^mu/hu^ KI mice. GAG values calculated include **(A)** Fold over TfR^mu/hu^ KI and **(B)** Percent reduction from vehicle treated *Ids* KO;TfR^mu/hu^ KI mice; n = 5 per group.

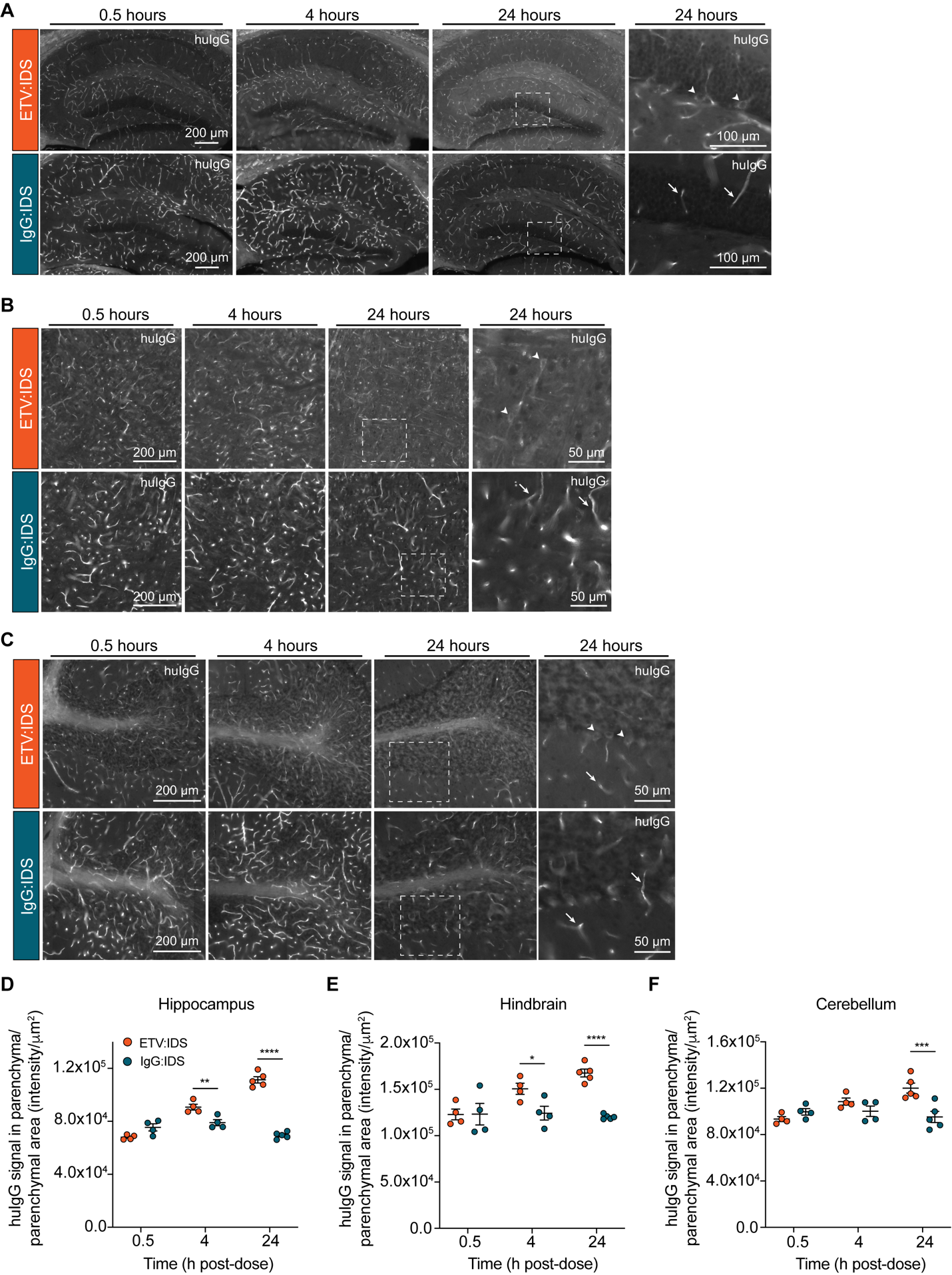

**Supplemental Figure 1.** **Immunohistochemical localization of ETV:IDS demonstrates more effective distribution into the brain parenchyma compared to IgG:IDS across multiple brain regions.** The distribution of ETV:IDS and IgG:IDS was assessed using sagittal brain sections from TfR^mu/hu^ KI mice at 0.5, 4, and 24 hours after an intravenous dose of 10 mg/kg. Schematic indicating approximate location of sagittal brain regions of interest (ROI) for images and quantification is found in Figure 4A. Sections were immunostained with antibodies against huIgG and imaged using a wide field fluorescence slide scanner in the **(A)** hippocampus, **(B)** hindbrain, and **(C)** cerebellum. Dashed boxes indicate regions shown at higher magnification displayed in the far-right panel. Arrows indicate huIgG staining localized to vascular profiles while arrowheads indicate cellular internalization of huIgG staining. Quantification of huIgG staining in the parenchyma of the **(D)** hippocampus, **(E)** hindbrain, and **(F)** cerebellum was calculated based on the total sum intensity of all parenchymal staining in the ROI divided by the total parenchymal area in the ROI. A custom macro script was used to identify blood vessels present in the tissue and masked out of subsequent image analyses; n = 4-5 per group. Graphs display mean ± SEM and p values: two-way ANOVA with Sidak’s multiple comparison test; * p ≤ 0.05, ** p ≤ 0.01, *** p ≤ 0.001 and **** p ≤ 0.0001.
